# A link between zinc uptake, bile salts, and a capsule required for virulence of a mastitis-associated extraintestinal pathogenic *Escherichia coli* strain

**DOI:** 10.1101/2020.06.25.172775

**Authors:** Michael A. Olson, Aleksander Grimsrud, Amanda C. Richards, Matthew A. Mulvey, Eric Wilson, David L. Erickson

## Abstract

Extraintestinal pathogenic *Escherichia coli* (ExPEC) are major causes of urinary and bloodstream infections. ExPEC reservoirs are not completely understood. Some mastitis-associated *E. coli* (MAEC) strains carry genes associated with ExPEC virulence, including metal scavenging, immune avoidance, and host attachment functions. In this study, we investigated the role of the high-affinity zinc uptake (*znuABC*) system in the MAEC strain M12. Elimination of *znuABC* moderately decreased fitness during mouse mammary gland infections. The Δ*znuABC* mutant strain exhibited an unexpected growth delay in the presence of bile salts, which was alleviated by the addition of excess zinc. We isolated Δ*znuABC* mutant suppressor mutants with improved growth of in bile salts, several of which no longer produced the K96 capsule made by strain M12. Addition of bile salts also reduced capsule production by strain M12 and ExPEC strain CP9, suggesting that capsule synthesis may be detrimental when bile salts are present. To better understand the role of the capsule, we compared the virulence of mastitis strain M12 with its unencapsulated Δ*kpsCS* mutant in two models of ExPEC disease. The wild type strain successfully colonized mouse bladders and kidneys and was highly virulent in intraperitoneal infections. Conversely, the Δ*kpsCS* mutant was unable to colonize kidneys and was unable to cause sepsis. These results demonstrate that some MAEC may be capable of causing human ExPEC illness. Virulence of strain M12 in these infections is dependent on its capsule. However, capsule may interfere with zinc homeostasis in the presence of bile salts while in the digestive tract.

## INTRODUCTION

*Escherichia coli* strains are abundant members of the healthy intestinal flora of most mammals and birds and may also be obligate or opportunistic pathogens. Strains that reside in the digestive tract but cause disease in other tissues are termed extraintestinal pathogenic *E. coli* (ExPEC) [1]. ExPEC are a major cause of urinary tract infections, neonatal meningitis, pneumonia, and sepsis in humans. ExPEC also cause several diseases of agricultural importance, including airway infections and septicemia in poultry [2]. Avian pathogenic strains present in poultry products represent a significant risk for human infection [3].

ExPEC strains are not derived from a single lineage, but rather arise from frequent recombination within diverse phylogenetic backgrounds [4]. Furthermore, there are no genes that are universally present in all ExPEC strains. Many specific genes have been shown to contribute to virulence of individual strains during experimental infections. Among these are genes that help the bacteria to attach to and invade their hosts, resist the antimicrobial effects of serum, scavenge for metal ions, or produce toxins [5].

*E. coli* are also the most frequent cause of bovine mastitis, which is responsible for billions of dollars in losses annually to dairy producers [6]. The strains that cause mastitis (Mastitis-Associated *E. coli* or MAEC) have traditionally been viewed as commensals that cause disease solely through triggering inflammation in the mammary gland [7]. As with ExPEC, no single gene is present in all MAEC. Several recent comparative genomic studies of MAEC have highlighted the diversity of these bacteria [8–12]. They have also noted that some MAEC carry genes associated with virulence of ExPEC strains [12–14]. The function of these ExPEC virulence genes in MAEC has not been investigated. As these genes are also commonly found in commensal strains that are not known to cause extraintestinal disease, their purpose may be to facilitate gastrointestinal colonization and persistence [15, 16]. These genes could also increase bacterial survival and tissue damage during mammary gland (MG) infection by some strains. Finally, it is also possible that some MAEC are fully capable of causing ExPEC-like disease, and pathogenesis in extraintestinal sites depends on these virulence-associated genes.

We previously conducted a genome-wide screen to identify fitness factors needed by a MAEC strain (M12) to grow in milk and colonize mouse MGs [12]. We demonstrated a role for an individual MAEC gene in mastitis for the first time. An M12 mutant lacking the *fecA* gene was unable to grow in milk and was attenuated during MG infection. The *fecA* gene encodes the ferric dicitrate receptor and is enriched among MAEC strains [17], but it is not known to contribute to other ExPEC disease. ExPEC-associated virulence genes were also identified as potentially contributing to MG colonization. These included genes for group III capsule biosynthesis identical to those of the K96 serotype [18]. Elimination of the *kpsCS* capsule genes from strain M12 resulted in a modest decrease in fitness in mouse MG colonization. The screen also suggested that the high affinity zinc uptake system encoded by the *znuABC* genes may be important in colonizing MGs.

Zinc is an essential cofactor for many enzymes and is essential for bacteria to overcome oxidative stress generated by innate immune responses. For example, superoxide dismutases (SODs) utilize different metals as cofactors, and zinc is a cofactor for SodC (CU/Zn-SOD). Neutrophils sequester zinc by releasing calprotectin during an active infection, thereby inhibiting SodC function [19, 20]. Infectious bacteria can scavenge for zinc using a variety of mechanisms, including high-affinity transport systems. The periplasmic zinc-binding protein ZnuA, integral membrane subunit ZnuB, and ATP-binding subunit ZnuC actively import scarce zinc ions across the inner membrane. These genes enhance virulence of several bacterial pathogens including ExPEC [21–26] but their role in MG colonization is unknown. High numbers of neutrophils infiltrate infected MGs during mastitis, so it is possible that expression of the Znu proteins benefit MAEC strains in this environment.

In this study, we investigated the role of the ExPEC-virulence associated *znuABC* genes in the M12 strain. We show that a M12 Δ*znuABC* mutant has decreased survival in mouse MGs. We also observed that the M12 Δ*znuABC* mutant has a pronounced growth delay when exposed to bile salts. Further examination of this growth defect uncovered a link between the presence of bile salts, zinc uptake and capsule production, where capsule appears to limit bacterial growth when bile salts are present. Conversely, we demonstrate that capsule synthesis is required for mastitis strain M12 to cause ExPEC-like disease including urinary tract infection and sepsis.

## RESULTS

### Role of *znuABC* in colonization of mouse mammary glands

Our previous TnSeq analysis suggested that the high-affinity zinc transport system (*znuABC*) is essential for MAEC strain M12 to colonize mouse MGs [12]. The *znuABC* locus contains two divergently transcribed units; *znuCB* are transcribed together while *znuA* is transcribed divergently (Fig. 1A). Reexamination of our TnSeq data suggested that Tn insertions in *znuA* or *znuC* but not in *znuB* are detrimental to M12 colonization of MGs [27]. In order to ascertain the importance of this system, we chose to delete the entire *znuABC* gene cluster from strain M12. To confirm that the Δ*znuABC* mutant strain was sensitive to zinc limiting conditions, the mutant and the wild type M12 strains were cultured in the presence of the divalent cation chelator EDTA. The Δ*znuABC* grew significantly more slowly compared to the wild type strain. Growth of the Δ*znuABC* mutant strain resembled that of the wild type when excess zinc was added to the media (Fig. 1B). We then tested the fitness of the mutant strain in conditions relevant to mastitis, including growth in milk and during infection of lactating mouse MGs. Competition assays demonstrated that loss of *znuABC* did not significantly affect fitness during growth in milk, but it resulted in an approximately 10-fold competitive disadvantage compared to wild type bacteria in mouse MGs (Fig. 1C).

**Figure 1.**
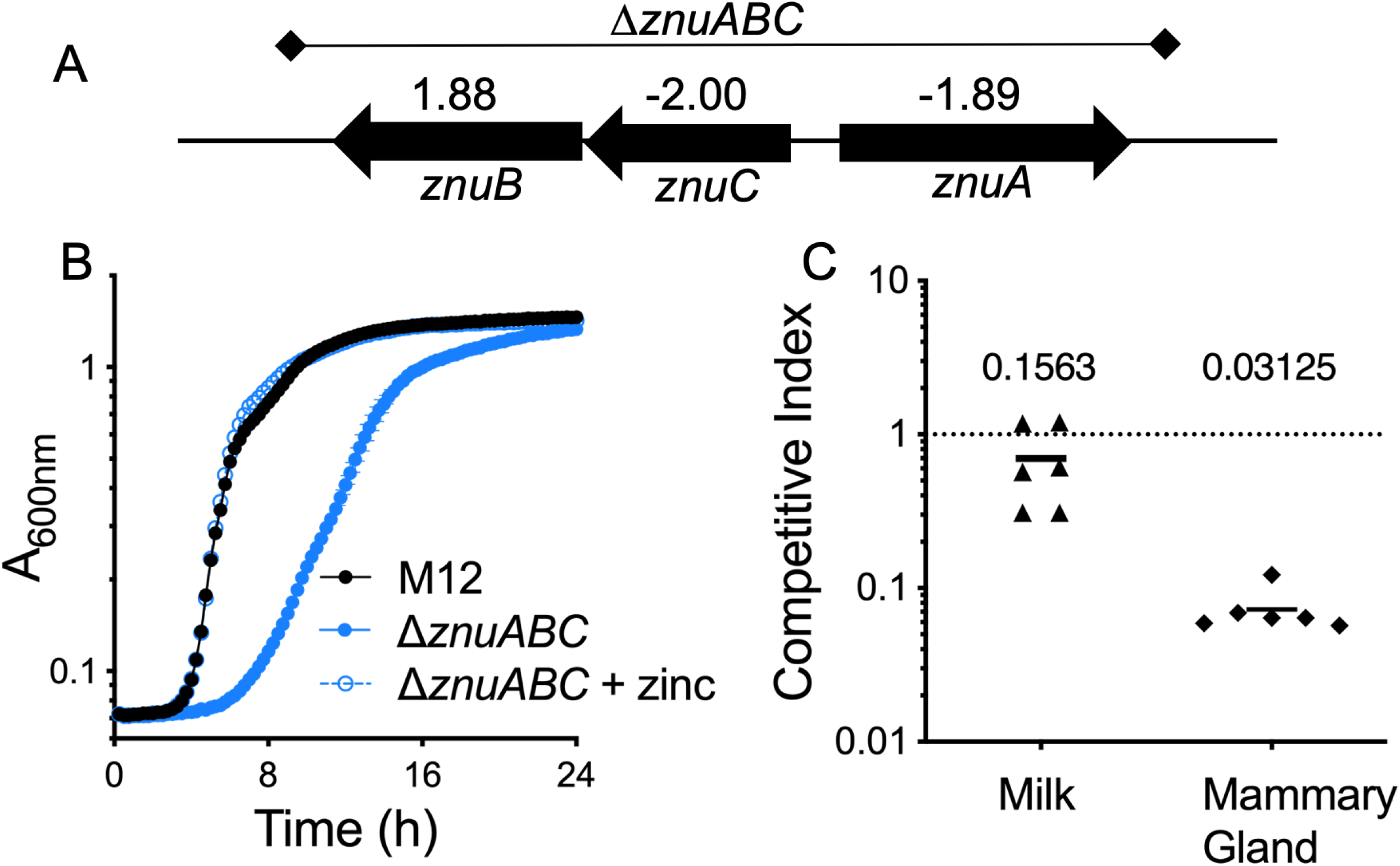
Role of the *znuABC* zinc transporter in the mastitis-associated *E. coli* M12 strain. (A) The *znu* locus of strain M12 and the 2560 bp region that was deleted to create the Δ*znuABC* mutant strain. The normalized TnSeq fitness scores predicted for each gene as calculated by Rendon et al. [27] are indicated. (B) Growth of M12 wild type and the Δ*znuABC* mutant in LB broth containing 0.5 mM EDTA. The mutant displayed a significant delay in growth that was fully restored with the addition of 100 μM zinc. (C) Competitive fitness of the Δ*znuABC* mutant compared with the M12 wild type strain during growth in milk or in mouse mammary glands. Equal ratios of both strains were inoculated in unpasteurized bovine milk or injected through the teat canal of lactating mice. Bacterial numbers were determined after 8 hours and 24 hours respectively. The mutant strain was significantly less fit (*p=0.03125 by Mann-Whitney test) than the wild-type strain in mammary glands but not in milk.

### Effect of *znuABC* on growth in the presence of bile salts

In these experiments, the competing strains were selected on MacConkey agar plates after growth in milk or mammary glands. We observed that when the Δ*znuABC* mutant was grown on MacConkey agar plates, colonies took 3-4 days to appear, but when grown on LB agar it grew at the same rate as the wild type strain. Bile salts and crystal violet are the primary constituents of MacConkey agar that select for Gram-negative enteric bacteria. In order to determine what was responsible for the delayed growth of the mutant strain, we tested growth in LB media containing bile salts or crystal violet. While crystal violet had no effect, the Δ*znuABC* mutant exhibited a delayed growth phenotype in media containing 2% bile salts (Fig. 2A). Total growth yield was the same for both wild type and mutant bacteria. The growth rate during exponential phase appeared to be the same, but the mutant did not enter exponential growth phase until 12-14 hours after inoculation.

**Figure 2.**
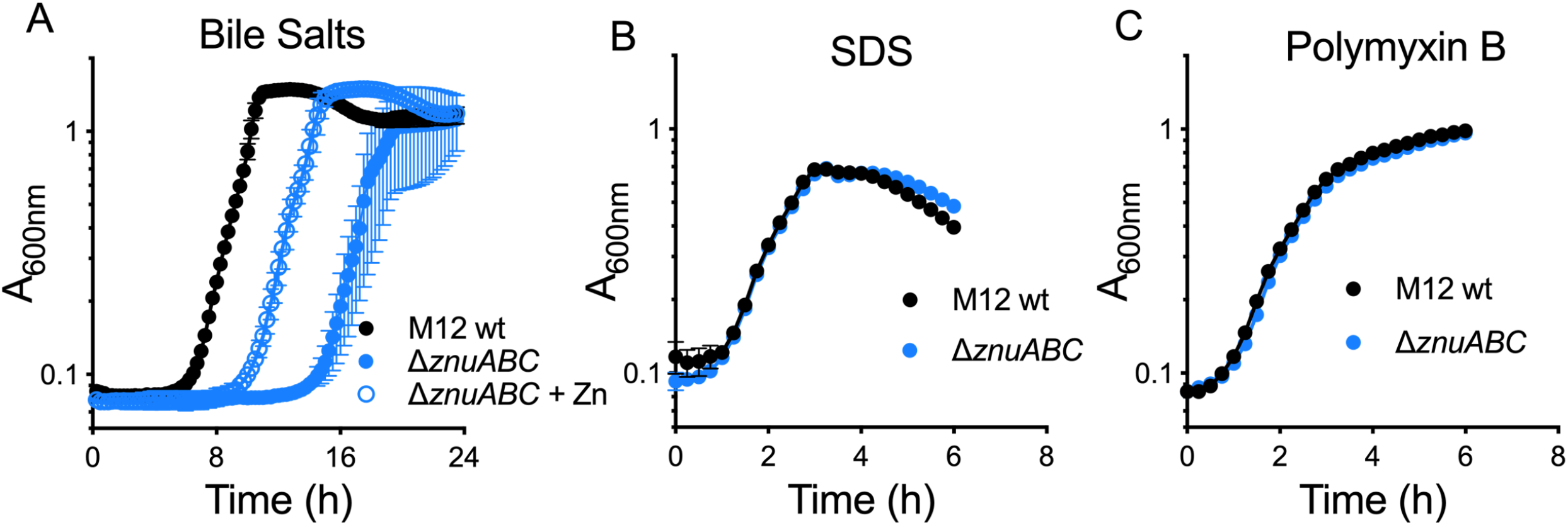
Loss of *znuABC* delays growth of strain M12 specifically in the presence of bile salts. (A) Growth of M12 or Δ*znuABC* mutant in LB media containing 2% bile salts. The mutant strain entered exponential growth phase several hours later than the wild type but had similar absorbance at 600 nm and during the saturation phase. Addition of 100 μM zinc to the media decreased the time it took for the Δ*znuABC* mutant to enter exponential phase. Identical growth curves of the wild type M12 and Δ*znuABC* mutant in 5% SDS (B) or 2 μg/ml of polymyxin E (C) showing that the mutation does not confer a generalized membrane defect.

Bile salts have detergent qualities that damage bacterial membranes. To determine if *ΔznuABC* had a generalized membrane defect, we investigated whether the Δ*znuABC* mutant had attenuated or delayed growth in SDS detergent or polymyxin B antibiotic, both of which can disrupt membrane integrity. The Δ*znuABC* did not have a growth defect in either SDS or polymyxin (Fig. 2B & C). As the Δ*znuABC* mutant does not have a generalized membrane defect, our results suggest a novel role for bile resistance that is dependent on zinc utilization. In support of this interpretation, addition of supplemental zinc (1 mM) to the media with bile salts decreased the time it took for the Δ*znuABC* mutant to enter exponential phase (Fig. 2A).

### Suppressor mutant screen suggests a link between K96 capsule and growth in bile salts

The Δ*znuABC* mutant grew to the same density as the wild type strain after 24 hours. We wanted to determine if these bacteria that had grown in the bile salt media would exhibit the same growth delay when subcultured in the same media. Transfer of these bacteria directly into LB+bile salts showed that they entered into exponential phase at the same time as the wild type strain. This suggested that the mutant strain adapted to the bile salts, either through changes in gene expression or through compensatory mutations or both. We then plated the bile-salt-adapted mutant strain on LB agar without bile salts to obtain single colonies. These colonies were then cultured in broth with bile salts (Fig. 3A) to see if the lag in growth reappeared. The “adapted” mutants grew at the same rate as the wild type strain, suggesting that suppressor mutants readily arose among the Δ*znuABC* mutant population in the presence of bile salts.

**Figure 3.**
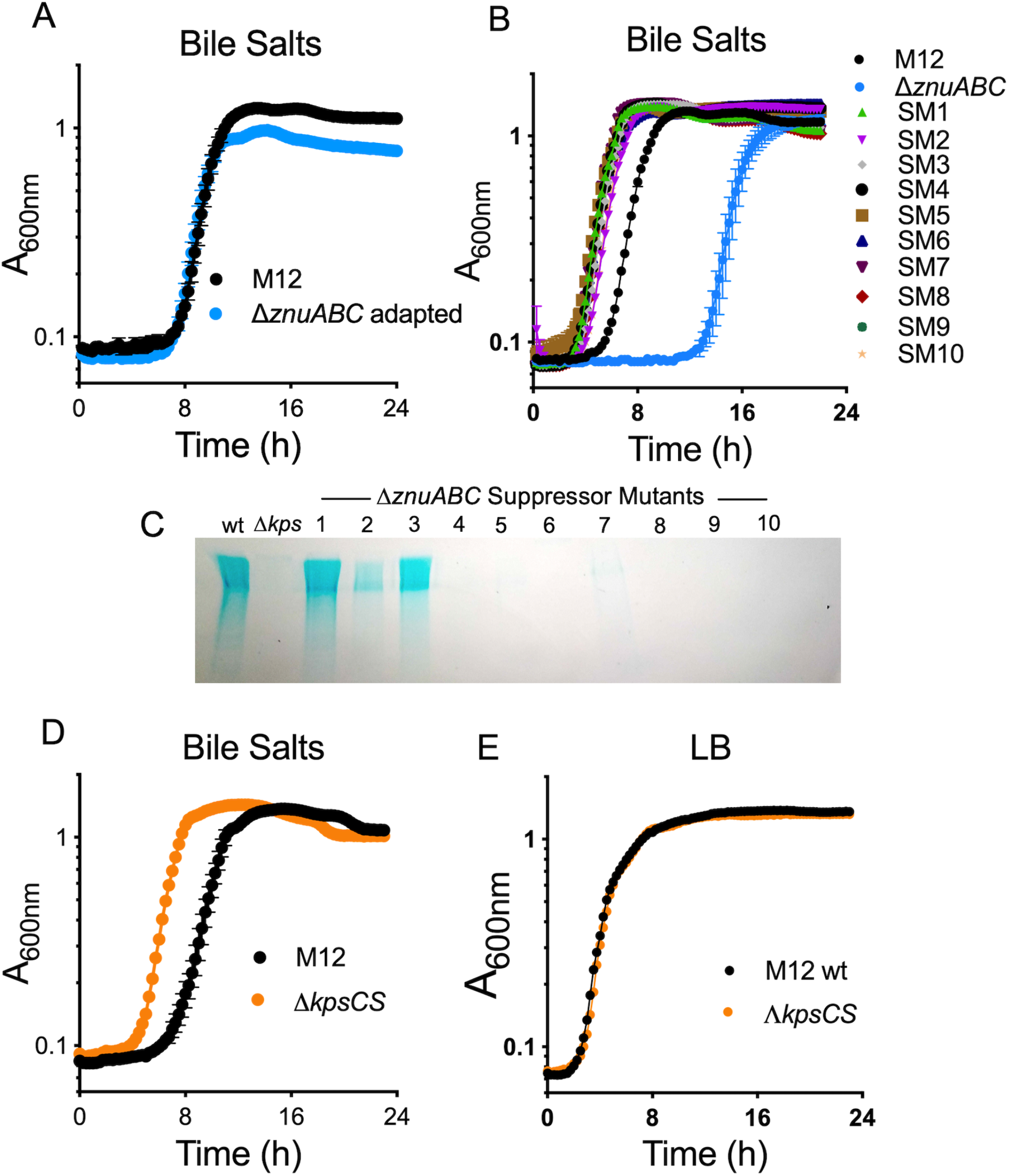
Suppressor mutations in the Δ*znuABC* background that restore growth in bile salts also eliminate capsule synthesis. (A) Saturated cultures of the wild type strain and the Δ*znuABC* mutant grown in LB + bile salts and subcultured in the same media exhibited similar growth rates, suggesting the possibility of suppressor mutations arising in the population. (B) Ten independently derived suppressor mutants had increased growth in LB media containing 2% bile salts when compared to the wild type strain. Whole-genome sequencing indicated that suppressor mutants 2, 8, and 10 contained SNPs that mapped to predicted capsule synthesis genes. (C) Gel electrophoresis and alcian blue staining shows that 8 of 10 suppressor mutants produce less capsule than the wild type strain. (D, E) A Δ*kpsSC* mutant unable to produce capsule reaches exponential phase faster than the wild type strain when grown in LB with bile salts but growth in LB is indistinguishable from the wild type strain.

To identify the genetic basis for these suppressor mutations, we serially passaged ten separate cultures derived from the Δ*znuABC* parent strain in the presence of 2% bile salts for approximately 200 generations. From these cultures, we isolated individual colonies on LB agar plates and then tested their growth profiles in bile salts compared to the wild type or Δ*znuABC* strains (Fig. 3B). All of the colonies we isolated grew more rapidly than the wild type strain in LB with bile salts. The genomes of the ten suppressor mutants as well as the Δ*znuABC* mutant parent strain were sequenced and non-synonymous SNPs within predicted coding regions were identified (Table 1). Three of the mutants had nucleotide substitutions in the *rpoA* gene encoding the alpha-subunit of RNA polymerase. All three mutations are predicted to change asparagine 294 to a histidine or lysine. Four mutants contained SNPs in the *dedD* or in *ftsK* genes that are predicted to result in non-functional alleles, either eliminating start codons or introducing premature stop codons. DedD and FtsK are both proteins involved in cell division. Finally, three suppressor mutants contained SNPs in glycosyltransferase genes found in a cluster associated with biosynthesis and export of a K96 group III capsule.

**Table 1.**
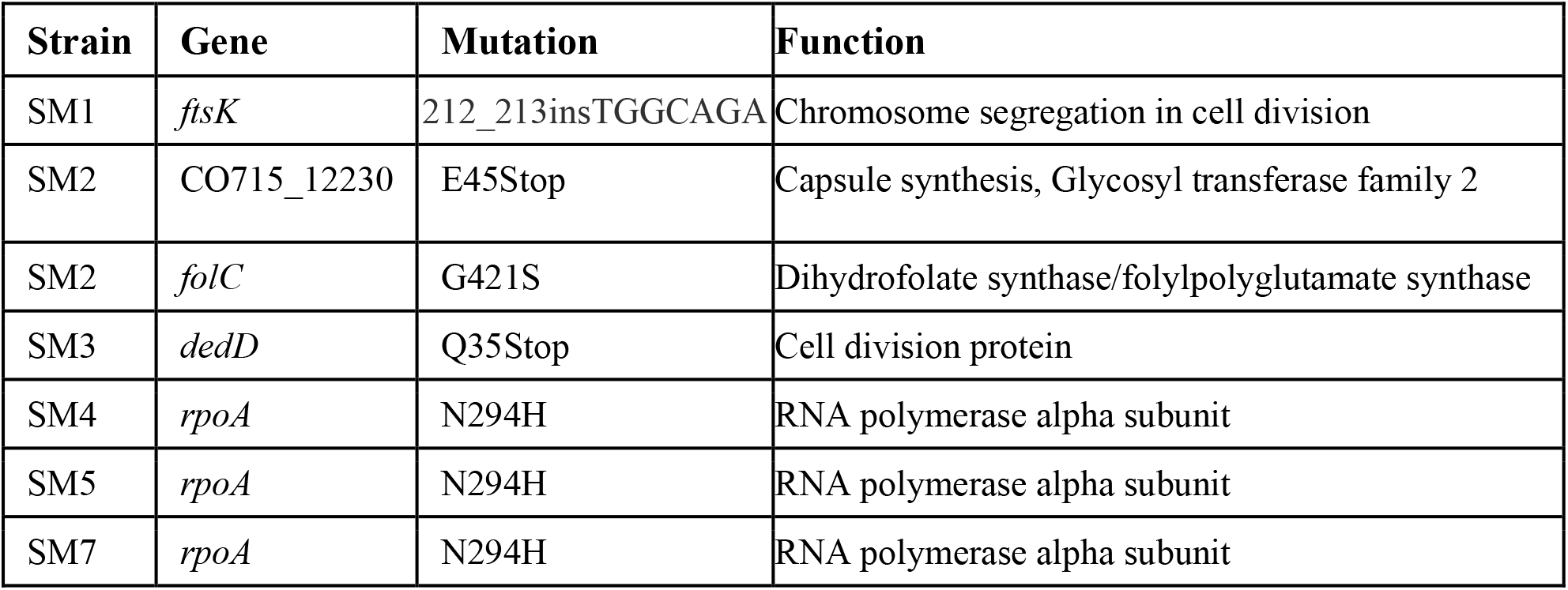
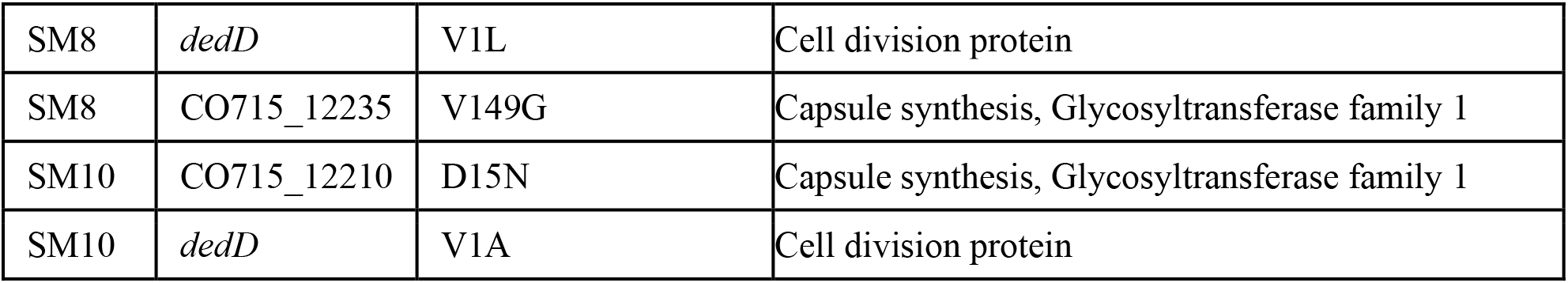
SNPs identified in Δ*znuABC* suppressor mutants with enhanced growth in bile salts.

| Strain | Gene | Mutation | Function |
| --- | --- | --- | --- |
| SM1 | *ftsK* | 212_213insTGGCAGA | Chromosome segregation in cell division |
| SM2 | CO715_12230 | E45Stop | Capsule synthesis, Glycosyl transferase family 2 |
| SM2 | *folC* | G421S | Dihydrofolate synthase/folylpolyglutamate synthase |
| SM3 | *dedD* | Q35Stop | Cell division protein |
| SM4 | *rpoA* | N294H | RNA polymerase alpha subunit |
| SM5 | *rpoA* | N294H | RNA polymerase alpha subunit |
| SM7 | *rpoA* | N294H | RNA polymerase alpha subunit |
| SM8 | *dedD* | V1L | Cell division protein |
| SM8 | CO715_12235 | V149G | Capsule synthesis, Glycosyltransferase family 1 |
| SM10 | CO715_12210 | D15N | Capsule synthesis, Glycosyltransferase family 1 |
| SM10 | *dedD* | V1A | Cell division protein |

These results suggested a link between capsule production and growth in bile salts. Therefore, we sought to determine whether these glycosyltransferase gene mutations altered capsule production. We also wanted to determine whether the other suppressor mutants produce capsule normally. We compared capsule synthesis of M12 wild type, the Δ*kpsCS* mutant, and each of the suppressor mutant strains (Fig. 3C) using SDS-PAGE and alcian blue staining. The suppressor mutants with changes in capsule glycosyltransferase genes did not produce detectable capsule. Surprisingly, several of the other suppressor mutants also failed to produce capsule, even though their mutations did not map to predicted capsule loci. This suggested that capsule production is detrimental to growth of strain M12 in bile salts. To test this directly, growth of the wild type M12 strain and the Δ*kpsCS* mutant were compared in LB or LB+2% bile salts (Fig. 3 D&E). Although growth in LB was identical, the Δ*kpsCS* mutant strain entered exponential phase more quickly than the wild type strain when bile salts were present.

Our results suggested that capsule may delay exponential growth of the bacteria when bile salts are present. This raised the possibility that capsule production may be influenced by the presence of bile salts. To test this, we grew the wild type strain M12 in LB or in LB+2% bile salts to examine how these conditions affected capsule synthesis. We also tested ExPEC strain CP9, which belongs to serotype K54. The chemical structure of K54 and K96 capsules are nearly identical [28]. When analyzed by alcian blue staining of gels, capsule synthesis appeared to be strongly repressed in both M12 and CP9 strains when grown in bile salts (Fig. 4A). We also used flow cytometry to measure capsule attached to intact bacteria. K54 antiserum bound strongly to both M12 and CP9 bacteria when grown in LB, but far more capsule-negative bacteria were detected when they were grown in LB + 2% bile salts (Fig. 4B).

**Figure 4.**
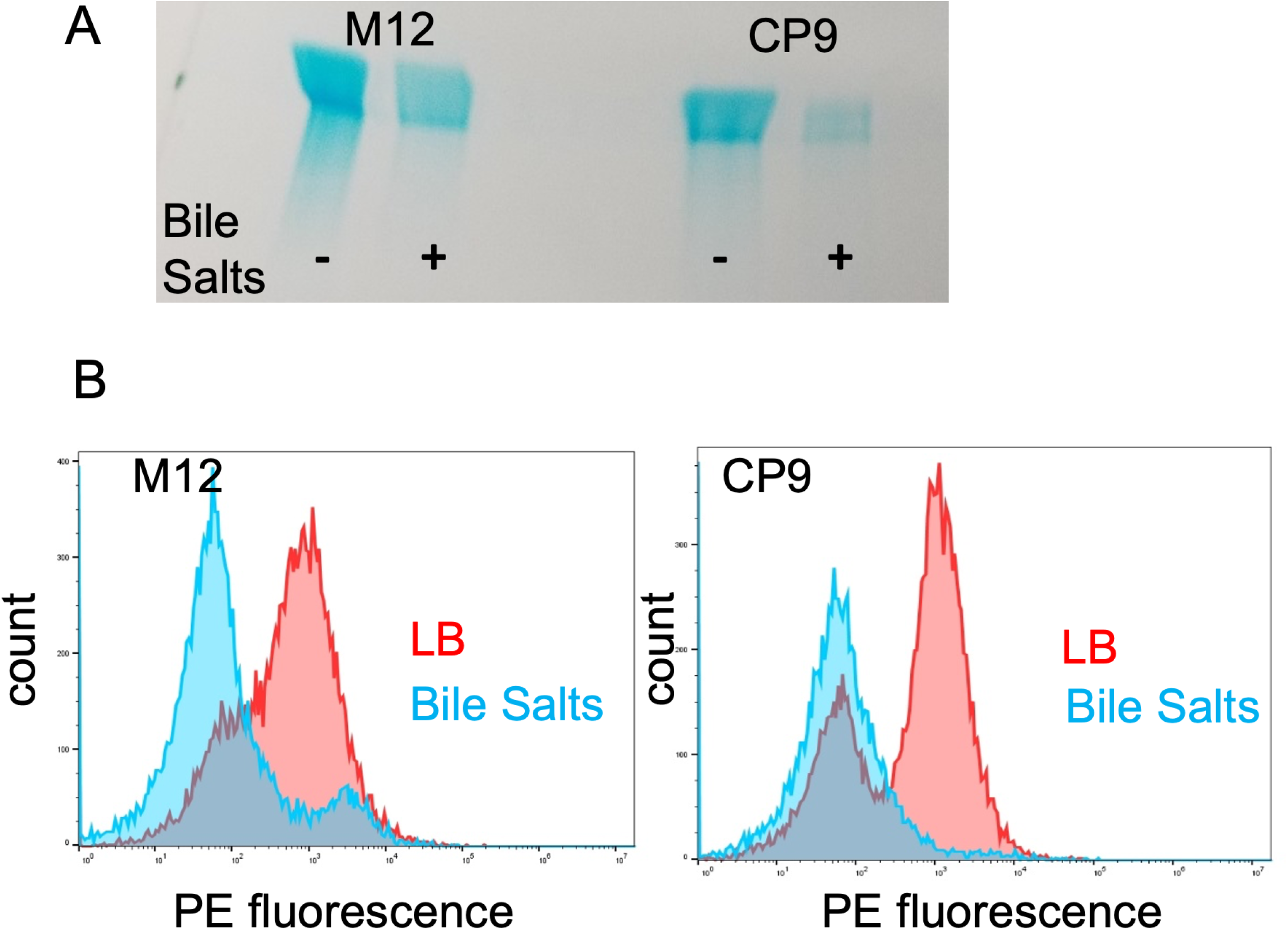
Growth in bile salts reduces capsule expression by strain M12 and ExPEC strain CP9. (A) Gel electrophoresis and alcian blue staining of capsule produced by M12 and CP9 grown in LB media or LB media containing 2% bile salts. (B) Capsule measurement by flow cytometry. In both M12 and CP9, the proportion of bacteria that were positively stained with K54 α-capsule serum detectable was less when grown in bile salts than in LB alone.

### Mastitis strain M12 causes urinary tract infection and sepsis in mice

MAEC strains circulate in the digestive tracts of dairy cattle and colonize mammary glands via environmental contamination. Our results suggest that capsule production may slow MAEC growth in the digestive tract where bile salts are present. Our previous work had also shown that the capsule of strain M12 contributes modestly to MG infection but is not required, which prompted us to further investigate the role of this capsule. Several MAEC strains possess similar genes for type II or type III capsules. They are required for some ExPEC strains to colonize the urinary tract and/or bloodstream. To investigate whether strain M12 could also infect these tissue sites, we used established rodent models of these human infections. First, we infected mice via intraperitoneal injection to mimic sepsis caused by ExPEC. We compared strain M12 with strain CP9 that is known to cause sepsis in this model. The wild type M12 strain was highly virulent in these mice, which exhibited signs of terminal illness and were euthanized after 24 h. High numbers of bacteria, similar to strain CP9, were recovered from infected spleens when the mice were sacrificed (Fig. 5A). We also tested whether the K96 capsule was important for strain M12 to cause sepsis. In contrast to the wild-type strain, mice infected with the M12 capsule mutant strain (Δ*kpsCS*) appeared healthy and far fewer bacteria were recovered from the spleens. We then tested whether strain M12 could infect the urinary tracts of mice. As with the intraperitoneal infections, strain M12 was very successful in the urinary tract, colonizing the bladders and kidneys of infected mice to high levels (Fig. 5B). The M12 Δ*kpsCS* mutant also efficiently colonized the bladders of the mice. However, the mutant strain was very attenuated in the kidneys and very few bacteria were recovered (Fig. 5B).

**Figure 5.**
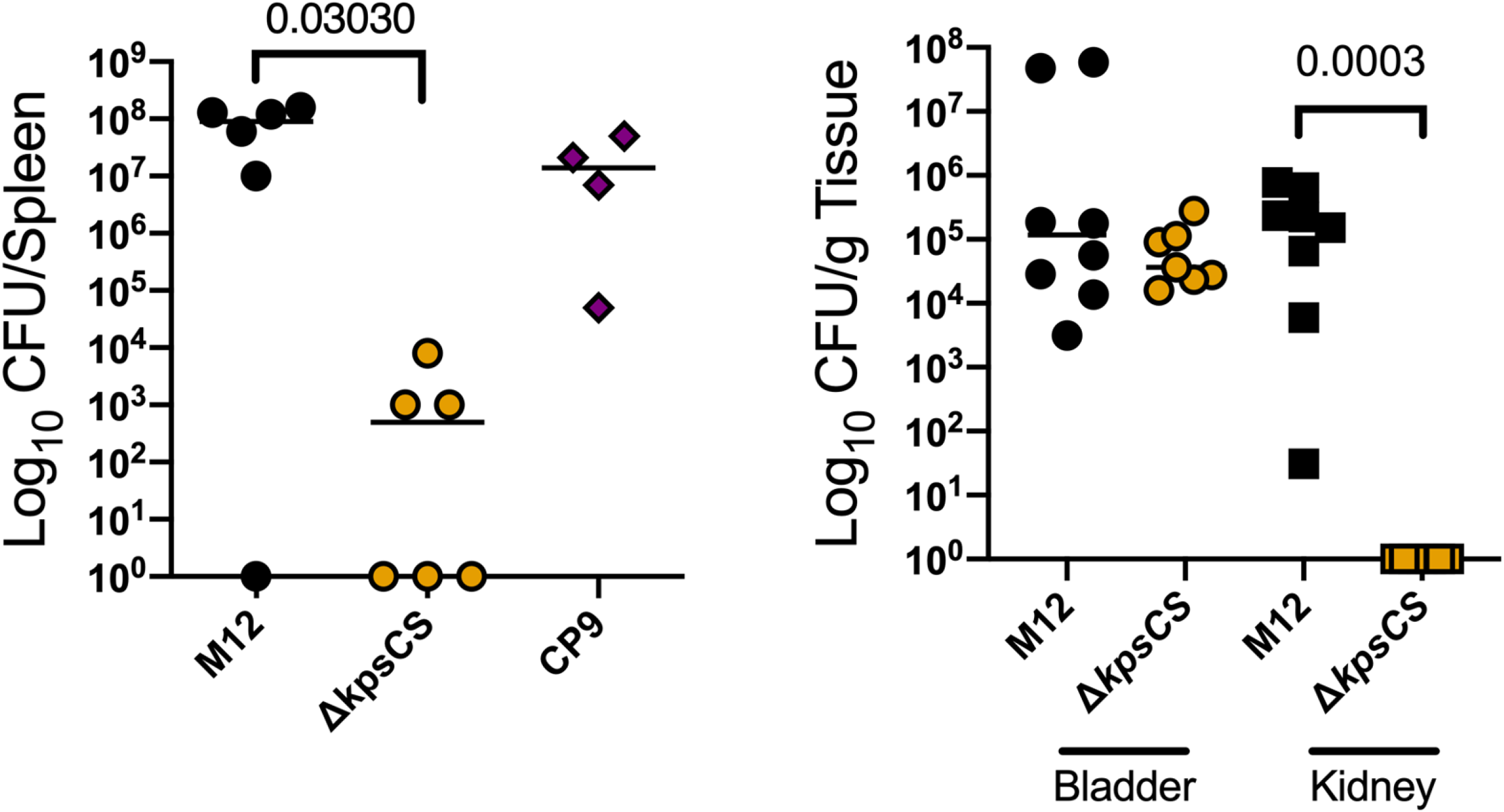
Virulence of mastitis-associated strain M12 in ExPEC infections depends on capsule production. (A) M12 wild type, M12 Δ*kpsCS* mutant and ExPEC CP9 strains (~5×10^5^ CFU) were injected into the intraperitoneal cavity of C57BL/6 mice and bacterial loads in the spleens were determined at 24 hours. Both M12 wild type and CP9 strains were recovered at high levels in the spleen while the Δ*kpsCS* mutant was severely attenuated (p=0.0303 by Mann-Whitney test). (B) Adult female Swiss-Webster mice were inoculated via transurethral catheterization and sacrificed 72 hours later. Both M12 wild type and Δ*kpsCS* colonized the bladders of the mice to similar levels but in the kidney the mutant strain was not detected (p=0.0003 by Mann-Whitney test).

## DISCUSSION

ExPEC are not defined by their lineage or carriage of any particular gene, but rather by their capacity to colonize specific tissue sites. However, some factors do tend to be more common in specific pathotypes including P fimbriae in uropathogenic strains [29], aerobactin siderophore and increased serum survival (*iss*) in avian pathogenic strains ([30]), and sialic acid capsule in neonatal meningitis *E. coli* strains [31]. MAEC strains have not previously been grouped under the ExPEC umbrella, even though some carry these virulence genes. Recent efforts to better understand the genomes of MAEC strains have uncovered certain features that tend to be associated with this group of bacteria [8, 9, 11, 32, 33]. For example, MAEC are more likely than other strains to express the ferric dicitrate receptor, and this improves bacterial growth in milk and lactating mammary glands [12]. It is now clear that all *E. coli* are not equally capable of causing mastitis and that the disease outcome depends greatly on the nature of the infecting strain rather than simply on host factors alone. For instance, strains that cause persistent mastitis tend to be more motile and more resistant to complement than those that cause transient disease [33]. Our results indicate that some classic ExPEC factors such as capsule and metal scavenging systems may also enhance growth of some strains in MGs. More frequent systemic spread or severe tissue damage would logically follow from prolific growth in the udder, and these ExPEC virulence factors may also contribute to bacterial survival beyond the MG.

Our interest in the role of the *znuABC* system stemmed from our previous TnSeq data that indicated it is required for M12 fitness in MGs and during *Galleria mellonella* infections [12], suggesting that the bacteria experience zinc starvation in these environments. Psoriasin (S100A7), a zinc-binding protein produced by keratinocytes, is found abundantly in mammary epithelia and is capable of inhibiting *E. coli* growth in vitro or in skin lesions [34, 35]. Neutrophils also employ zinc sequestration as an antimicrobial strategy. They infiltrate infected MGs quickly after infection and are critical to the containment of the bacteria and limiting tissue damage [36]. They release calprotectin to significantly limit extracellular zinc. Once inside phagosomes, the bacteria also face toxic levels of excess zinc. Thus, systems to maintain homeostasis may be key fitness determinants.

We were surprised to find that bile salts delayed growth of the M12 Δ*znuABC* mutant. Mutants lacking these genes have been made in other *E. coli* strain backgrounds but no similar growth defects on MacConkey agar were reported. Whether *znuABC* mutations in other strain backgrounds confer growth defects in bile salts deserves further investigation. Bile concentrations reach levels between 0.2–2.0% in the small intestine [37]. Due to the detergent properties of bile, it is capable of disrupting bacterial membranes and crossing into the cell, which can induce DNA damage and oxidative stress [38, 39]. Bacteria employ multiple tactics to combat the disruptive effects of bile, including outer membrane modifications, efflux pumps, and DNA repair systems [37]. It is likely that variation of cell envelope features such as capsule or lipopolysaccharide greatly influence the sensitivity of different strains to bile salt stress. This is supported by our finding that loss of capsule enabled the M12 Δ*znuABC* mutant to grow in the presence of bile salts. The K96 capsule made by M12 consists of glucuronic acid and rhamnose and has a strong negative charge [28]. It is conceivable that this capsule interferes with zinc cation movement into the cells when the high affinity Znu system is not present and zinc is not readily available. Our results indicate that bile salts may confer this type of stress, reducing the amount of zinc available for bacterial growth.

To our knowledge, there are no reports of bile salts specifically conferring zinc stress in other bacteria. However, the secondary bile acid deoxycholic acid can form organometallic complexes with zinc and other heavy metals which may make them unavailable for import, potentially affecting gene expression in many enteric bacteria [40–43]. *E. coli* O157:H7 is reported to increase expression of iron acquisition genes when exposed to bile, but repress key virulence genes including shiga toxin and the locus of enterocyte effacement [44]. We found that both the MAEC strain M12 and the human sepsis strain CP9 repress production of capsule in the presence of bile salts. These bacteria may benefit from repressing virulence factors while transiting through the small intestine until the appropriate target sites are reached; the colon in the case of EHEC, and extraintestinal tissues for M12 and CP9. The relevant sensing mechanisms and the level at which the regulation is achieved remain to be identified. Furthermore, it is not yet clear whether the influence of bile salts on capsule production occurs through its effect on zinc availability or via sensing bile salts directly. Zinc regulation of capsule production in bacteria is not without precedent; in *Streptococcus pyogenes*, high levels of zinc such as are found within neutrophil phagosomes inhibit the phosphoglucomutase enzyme needed to initiate capsule synthesis [45].

ExPEC capsules impact pathogenesis in non-uniform ways. More than 100 distinct capsule types have been described, which are categorized into four groups based on gene organization and mode of assembly and export [46]. Strain CP9 produces a K54 capsule belonging to group 3 that has a strong effect on virulence. CP9 mutants lacking this capsule were highly attenuated in mouse bloodstream infections and in a rat model of abscess formation [47]. However, loss of the K54 capsule did not have a measurable effect in the ability of strain CP9 to colonize mouse bladders or kidneys during urinary tract infections [48]. Most uropathogenic ExPEC strains produce capsule belonging to group 2. For instance, strain CFT073 produces a K2 capsule that was shown to have a modest effect in promoting colonization in mouse bladders [49, 50] and a more important role in kidneys [49]. In UPEC strain 536, production of the K15 capsule was proven to contribute dramatically to urovirulence in neonatal mice [51]. In these studies, mortality of the mice was measured and not bacterial loads in the bladder or kidneys, so it is unclear at which stage colonization was affected.

We have shown that an MAEC strain M12 colonizes both bladders and kidneys during experimental urinary tract infections and causes lethal sepsis in mice (Fig. 5). In both cases, the production of K96 capsule was a critical virulence determinant for colonization of kidneys and spleens. Previously our group showed that this mutant is also highly attenuated in *G. mellonella* [12]. It is interesting that the capsules of strains M12 and CP9 are identical but in CP9 the capsule is dispensable for kidney infection via the urinary tract [48], whereas our results show it is required in strain M12. Absence of the K96 capsule of M12 did not detectably alter bladder colonization (Fig. 5) and has a moderate effect in MGs. The exact function of the M12 capsule in these specific tissues is not yet understood. ExPEC capsules may have a role in avoiding killing by phagocytes [52]. The capsules of some ExPEC strains are required for serum resistance, while in other strains the loss of capsule has no effect [53–56]. ExPEC capsule can also help in the establishment of intracellular bacterial communities within epithelial cells of the urinary tract [57]. The fact that some MAEC strains carry capsule and other ExPEC virulence genes has been noted previously, but ours is the first study to show that a MAEC strain is capable of causing urinary tract infection and sepsis in established models of human disease. Non-pathogens may carry ExPEC virulence factors because they promote intestinal colonization rather than infection, depending on the strain background. However, as we have shown here, some MAEC strains may not be purely commensals or accidental pathogens, but rather versatile organisms fully capable of causing human illness in addition to bovine mastitis.

## EXPERIMENTAL PROCEDURES

### Bacterial strains and growth conditions

*E. coli* strain CP9 was generously supplied by xx. *E. coli* strain M12 and the M12 Δ*kpsCS* mutant were previously characterized [12]. *E. coli* strains were routinely grown in Luria-Bertani (LB) medium at 37°C. To select for mutant strains or carriage of plasmids, growth medium was supplemented with chloramphenicol (10 μg/ml), or ampicillin (100 μg/ml) as appropriate. For milk cultures, whole unpasteurized cow’s milk was obtained from a local supplier and used immediately or stored at −80°C until use. Growth curves were generated using a Bioscreen C system (Labsystems Oy, Helsinki, Finland). Bacterial cultures were standardized to 5×10^4^ CFU/mL and absorbance at 600 nm was measured every 15 minutes while continuously shaken at 37°C. LB growth medium was supplemented with 2% bile salts, 2 μg/ml polymyxin B, 5% SDS, 100 μM EDTA or 100 μM zinc chloride as appropriate. All chemicals were purchased from Sigma-Aldrich. For competition assays during growth in milk, equal ratios of M12 and Δ*znuABC* mutant was inoculated into whole unpasteurized cow’s milk and incubated at 37°C for 8 h. CFU ratios of the input and output for M12 and Δ*znuABC*, determined by plate counts, were used to calculate a competition index.

### Deletion of *znuABC*

Mutation of *znuABC* was performed via lambda red recombination in strain M12 carrying the plasmid pKD46 [58]. A PCR product was created using pRE112 [59] as a template to amplify the chloramphenicol acetyltransferase gene, with 50 bp overhangs homologous to the 5’ end of *znuB* or the 3’ end of *znuA*. Bacteria expressing the recombinase enzymes were electroporated with 500 ng of purified DNA, and potential mutants were selected by plating on LB agar containing chloramphenicol. Potential mutants were then screened by PCR using primers flanking the recombination site and also with primers internal to the *znuB* and *znuA* genes. The correct Δ*znuABC* mutation and absence of other SNPs was also confirmed by whole genome sequencing (Microbial Genome Sequencing Center, University of Pittsburgh).

### Suppressor mutant sequencing, assembly, and annotation

Total DNA was isolated from suppressor mutants using a ZR Fungal/Bacterial DNA MiniPrep kit (Zymoresearch). DNA sequencing libraries were prepared using the Illumina Nextera DNA library Prep kit with modifications [60]. DNA libraries were sequenced by Genewiz, Inc. (South Plainfield, NJ) and Illumina paired-end reads of 150 bp were generated on MiSeq version 2 sequencer. Contigs, annotations and SNP analyses were compiled using EnteroBase [61, 62].

### Capsule isolation and staining

Capsule production was visualized as previously described [63]. Briefly, bacteria were pelleted from 1 ml of saturated overnight cultures grown in LB broth. The bacterial cells were then resuspended in 50 μl of PBS and heated to 55°C for 30 min. The capsule material analyzed by electrophoresis on 10% SDS-polyacrylamide gels and staining with 0.125% alcian blue dye in 40% ethanol–5% acetic acid.

### Flow cytometry

Bacteria were grown in LB broth or LB broth supplemented with 2% bile salts for 24 h. Saturated overnight cultures were diluted 1:100 in 0.1% BSA-PBS with undiluted anti-K54 rabbit antisera (SSI Diagnostica) and incubated for 30 minutes on ice. Samples were washed with 0.1% BSA-PBS and stained with Goat anti-Rabbit IgG (H+L) secondary antibody, Alexa Fluor Plus 594 (Invitrogen) in the dark for 30 minutes. The fluorescence of individual bacterial cells was measured using a BD Accuri C6 flow cytometer and histograms were generated with FlowJo™ Software. Negative control samples contained bacteria with the secondary Ab only or without primary and secondary Ab.

### Ethics statement

Mouse experiments were performed in accordance with the recommendations found in the Guide for the Care and Use of Laboratory Animals of the National Institutes of Health (83). The protocol was reviewed and approved by the Institutional Animal Care and Use Committees of Brigham Young University and the University of Utah.

### Mouse infections

#### Mouse mammary gland infections

Lactating C57BL/6 mice between 9 and 12 weeks of age and 10 to 11 days postpartum were infected as previously described [12]. Briefly, a 50 μl volume of bacteria containing 250 CFU of both wild type M12 and the Δ*znuABC* mutant strain in PBS was injected directly through the teat canal into the ductal network of the 4^th^ left and 4^th^ right mammary glands of 3 mice using a 33-gauge needle with a beveled end. Pups were removed for 1 to 2 h after injections and then were returned. Mice were euthanized after 24 h and mammary gland tissue was harvested. Bacterial loads were determined by homogenizing tissue in 1 ml of PBS, performing serial dilutions and plating on selective agar. CFU ratios of the input and output for M12 and Δ*znuABC*, determined by plate counts, were used to calculate a competition index.

#### Mouse intraperitoneal infections

Equal numbers of male and female C57BL/6 mice between 9 and 12 weeks of age were used for all infections. Bacteria were grown overnight in LB, subcultured to an absorbance of 1.0 at 600 nn, and serially diluted in PBS. A 200 μl volume of bacteria containing 5×10^5^ CFU in PBS was injected directly into the intraperitoneal cavity using a 27 ½-gauge needle. The concentration of each inoculum was determined by serial dilution and colony counting after 24 h growth on LB agar plates. Mice were monitored for 24 h and spleens were harvested. Bacterial loads were determined by homogenizing entire spleens in 1 ml of PBS, performing serial dilutions, and colony counts.

#### Mouse urinary tract infections

Adult (~8 weeks) female Swiss-Webster mice (Charles River) were used in all experiments Bacteria were grown statically from frozen stocks in 20 mL M9 minimum medium for 24 h at 37°C. Bacteria were pelleted at 8,000 RCF for 8 minutes and resuspended in 3 mL sterile PBS. Mice were inoculated with a 50μl volume (5×10^8^ CFU) via transurethral catheterization and sacrificed 3 days later. Kidneys and bladders were weighed, homogenized, and CFU determined by serial dilution and plating.

### Statistical analyses

All mouse infection data were analyzed using GraphPad Prism 5.0. The statistical tests performed as well as the significance values are indicated in the individual figure legends.

## REFERENCES

1. Manges AR, Johnson JR. Reservoirs of Extraintestinal Pathogenic Escherichia coli. Microbiol Spectr. 2015;3(5). Epub 2015/11/07. doi: 10.1128/microbiolspec.UTI-0006-2012. PubMed PMID: 26542041.

2. Russo TA, Johnson JR. Proposal for a new inclusive designation for extraintestinal pathogenic isolates of Escherichia coli: ExPEC. J Infect Dis. 2000;181(5):1753–4. Epub 2000/05/24. doi: 10.1086/315418. PubMed PMID: 10823778.

3. Mellata M, Johnson JR, Curtiss R, 3rd. Escherichia coli isolates from commercial chicken meat and eggs cause sepsis, meningitis and urinary tract infection in rodent models of human infections. Zoonoses Public Health. 2017. doi: 10.1111/zph.12376. PubMed PMID: 28703468.

4. Johnson JR, Russo TA. Molecular Epidemiology of Extraintestinal Pathogenic Escherichia coli. EcoSal Plus. 2018;8(1). Epub 2018/04/19. doi: 10.1128/ecosalplus.ESP-0004-2017. PubMed PMID: 29667573.

5. Wiles TJ, Kulesus RR, Mulvey MA. Origins and virulence mechanisms of uropathogenic Escherichia coli. Exp Mol Pathol. 2008;85(1):11–9. Epub 2008/05/17. doi: 10.1016/j.yexmp.2008.03.007. PubMed PMID: 18482721; PubMed Central PMCID: PMCPMC2595135.

6. Blum SE, Heller ED, Leitner G. Long term effects of Escherichia coli mastitis. Vet J. 2014;201(1):72–7. doi: 10.1016/j.tvjl.2014.04.008. PubMed PMID: 24906501.

7. Suojala L, Pohjanvirta T, Simojoki H, Myllyniemi AL, Pitkala A, Pelkonen S, et al. Phylogeny, virulence factors and antimicrobial susceptibility of Escherichia coli isolated in clinical bovine mastitis. Vet Microbiol. 2011;147(3-4):383–8. Epub 2010/08/24. doi: 10.1016/j.vetmic.2010.07.011. PubMed PMID: 20729012.

8. Kempf F, Slugocki C, Blum SE, Leitner G, Germon P. Genomic Comparative Study of Bovine Mastitis Escherichia coli. PLoS ONE. 2016;11(1):e0147954. doi: 10.1371/journal.pone.0147954. PubMed PMID: 26809117; PubMed Central PMCID: PMCPMC4725725.

9. Leimbach A, Poehlein A, Vollmers J, Gorlich D, Daniel R, Dobrindt U. No evidence for a bovine mastitis Escherichia coli pathotype. BMC Genomics. 2017;18:359.

10. Richards VP, Lefebure T, Pavinski Bitar PD, Dogan B, Simpson KW, Schukken YH, et al. Genome based phylogeny and comparative genomic analysis of intra-mammary pathogenic Escherichia coli. PLoS ONE. 2015;10(3):e0119799. doi: 10.1371/journal.pone.0119799. PubMed PMID: 25807497; PubMed Central PMCID: PMCPMC4373696.

11. Goldstone RJ, Harris S, Smith DG. Genomic content typifying a prevalent clade of bovine mastitis-associated Escherichia coli. Scientific reports. 2016;6:30115. doi: 10.1038/srep30115. PubMed PMID: 27436046; PubMed Central PMCID: PMCPMC4951805.

12. Olson MA, Siebach TW, Griffitts JS, Wilson E, Erickson DL. Genome-Wide Identification of Fitness Factors in Mastitis-Associated Escherichia coli. Appl Environ Microbiol. 2018;84(2):e02190–17. Epub 2017/11/05. doi: 10.1128/AEM.02190-17. PubMed PMID: 29101196; PubMed Central PMCID: PMCPMC5752858.

13. Nuesch-Inderbinen M, Kappeli N, Morach M, Eicher C, Corti S, Stephan R. Molecular types, virulence profiles and antimicrobial resistance of Escherichia coli causing bovine mastitis. Vet Rec Open. 2019;6(1):e000369. Epub 2020/01/04. doi: 10.1136/vetreco-2019-000369. PubMed PMID: 31897302; PubMed Central PMCID: PMCPMC6924703.

14. Blum SE, Leitner G. Genotyping and virulence factors assessment of bovine mastitis Escherichia coli. Vet Microbiol. 2013;163(3-4):305–12. doi: 10.1016/j.vetmic.2012.12.037. PubMed PMID: 23374653.

15. Diard M, Garry L, Selva M, Mosser T, Denamur E, Matic I. Pathogenicity-associated islands in extraintestinal pathogenic Escherichia coli are fitness elements involved in intestinal colonization. J Bacteriol. 2010;192(19):4885–93. Epub 2010/07/27. doi: 10.1128/JB.00804-10. PubMed PMID: 20656906; PubMed Central PMCID: PMCPMC2944530.

16. Russell CW, Fleming BA, Jost CA, Tran A, Stenquist AT, Wambaugh MA, et al. Context-Dependent Requirements for FimH and Other Canonical Virulence Factors in Gut Colonization by Extraintestinal Pathogenic Escherichia coli. Infect Immun. 2018;86(3). Epub 2018/01/10. doi: 10.1128/IAI.00746-17. PubMed PMID: 29311232; PubMed Central PMCID: PMCPMC5820936.

17. Lin J, Hogan JS, Smith KL. Antigenic homology of the inducible ferric citrate receptor (FecA) of coliform bacteria isolated from herds with naturally occurring bovine intramammary infections. Clin Diagn Lab Immunol. 1999;6(6):966–9. PubMed PMID: 10548594; PubMed Central PMCID: PMCPMC95806.

18. Yang S, Xi D, Jing F, Kong D, Wu J, Feng L, et al. Genetic diversity of K-antigen gene clusters of Escherichia coli and their molecular typing using a suspension array. Can J Microbiol. 2018;64(4):231–41. Epub 2018/01/23. doi: 10.1139/cjm-2017-0620. PubMed PMID: 29357266.

19. Urban CF, Ermert D, Schmid M, Abu-Abed U, Goosmann C, Nacken W, et al. Neutrophil Extracellular Traps Contain Calprotectin, a Cytosolic Protein Complex Involved in Host Defense against Candida albicans. PLoS Pathog. 2009;5(10). doi: 10.1371/journal.ppat.1000639. PubMed PMID: WOS:000272033300043.

20. Sohnle PG, Collinslech C, Wiessner JH. THE ZINC-REVERSIBLE ANTIMICROBIAL ACTIVITY OF NEUTROPHIL LYSATES AND ABSCESS FLUID SUPERNATANTS. Journal of Infectious Diseases. 1991;164(1):137–42. doi: 10.1093/infdis/164.1.137. PubMed PMID: WOS:A1991FT63000020.

21. Gunasekera TS, Herre AH, Crowder MW. Absence of ZnuABC-mediated zinc uptake affects virulence-associated phenotypes of uropathogenic Escherichia coli CFT073 under Zn(II)-depleted conditions. Fems Microbiology Letters. 2009;300(1):36–41. doi: 10.1111/j.1574-6968.2009.01762.x. PubMed PMID: WOS:000270434400005.

22. Campoy S, Jara M, Busquets N, de Rozas AMP, Badiola I, Barbe J. Role of the high-affinity zinc uptake znuABC system in Salmonella enterica serovar Typhimurium virulence. Infect Immun. 2002;70(8):4721–5. doi: 10.1128/iai.70.8.4721-4725.2002. PubMed PMID: WOS:000176909400089.

23. Corbett D, Wang JH, Schuler S, Lopez-Castejon G, Glenn S, Brough D, et al. Two Zinc Uptake Systems Contribute to the Full Virulence of Listeria monocytogenes during Growth In Vitro and In Vivo. Infect Immun. 2012;80(1):14–21. doi: 10.1128/iai.05904-11. PubMed PMID: WOS:000298402500002.

24. D’Orazio M, Mastropasqua MC, Cerasi M, Pacello F, Consalvo A, Chirullo B, et al. The capability of Pseudomonas aeruginosa to recruit zinc under conditions of limited metal availability is affected by inactivation of the ZnuABC transporter. Metallomics. 2015;7(6):1023–35. doi: 10.1039/c5mt00017c. PubMed PMID: WOS:000356058300011.

25. Hood MI, Mortensen BL, Moore JL, Zhang YF, Kehl-Fie TE, Sugitani N, et al. Identification of an Acinetobacter baumannii Zinc Acquisition System that Facilitates Resistance to Calprotectin-mediated Zinc Sequestration. PLoS Pathog. 2012;8(12). doi: 10.1371/journal.ppat.1003068. PubMed PMID: WOS:000312907100023.

26. Sabri M, Houle S, Dozois CM. Roles of the extraintestinal pathogenic Escherichia coli ZnuACB and ZupT zinc transporters during urinary tract infection. Infect Immun. 2009;77(3):1155–64. Epub 2008/12/24. doi: 10.1128/IAI.01082-08. PubMed PMID: 19103764; PubMed Central PMCID: PMCPMC2643633.

27. Rendon JM, Lang B, Tartaglia GG, Burgas MT. BacFITBase: a database to assess the relevance of bacterial genes during host infection. Nucleic Acids Res. 2020;48(D1):D511–D6. Epub 2019/10/31. doi: 10.1093/nar/gkz931. PubMed PMID: 31665505; PubMed Central PMCID: PMCPMC7145566.

28. Jann B, Kochanowski H, Jann K. Structure of the capsular K96 polysaccharide (K96 antigen) from Escherichia coli O77:K96:H- and comparison with the capsular K54 polysaccharide (K54 antigen) from Escherichia coli O6:K54:H10. Carbohydr Res. 1994;253:323–7. Epub 1994/02/03. doi: 10.1016/0008-6215(94)80080-4. PubMed PMID: 8156556.

29. Bergsten G, Wullt B, Svanborg C. Escherichia coli, fimbriae, bacterial persistence and host response induction in the human urinary tract. Int J Med Microbiol. 2005;295(6-7):487–502. Epub 2005/10/22. doi: 10.1016/j.ijmm.2005.07.008. PubMed PMID: 16238023.

30. Johnson TJ, Giddings CW, Horne SM, Gibbs PS, Wooley RE, Skyberg J, et al. Location of increased serum survival gene and selected virulence traits on a conjugative R plasmid in an avian Escherichia coli isolate. Avian diseases. 2002;46(2):342–52. Epub 2002/06/14. doi: 10.1637/0005-2086(2002)046[0342:LOISSG]2.0.CO;2. PubMed PMID: 12061643.

31. Wijetunge DSS, Gongati S, DebRoy C, Kim KS, Couraud PO, Romero IA, et al. Characterizing the pathotype of neonatal meningitis causing Escherichia coli (NMEC). Bmc Microbiology. 2015;15. doi: 10.1186/s12866-015-0547-9. PubMed PMID: WOS:000362707400001.

32. Blum S, Sela N, Heller ED, Sela S, Leitner G. Genome analysis of bovine-mastitis-associated Escherichia coli O32:H37 strain P4. J Bacteriol. 2012;194(14):3732. doi: 10.1128/JB.00535-12. PubMed PMID: 22740662; PubMed Central PMCID: PMCPMC3393488.

33. Lippolis JD, Holman DB, Brunelle BW, Thacker TC, Bearson BL, Reinhardt TA, et al. Genomic and Transcriptomic Analysis of Escherichia coli Strains Associated with Persistent and Transient Bovine Mastitis and the Role of Colanic Acid. Infect Immun. 2018;86(1). Epub 2017/10/25. doi: 10.1128/IAI.00566-17. PubMed PMID: 29061709; PubMed Central PMCID: PMCPMC5736815.

34. Zhang GW, Lai SJ, Yoshimura Y, Isobe N. Messenger RNA expression and immunolocalization of psoriasin in the goat mammary gland and its milk concentration after an intramammary infusion of lipopolysaccharide. Vet J. 2014;202(1):89–93. Epub 2014/07/16. doi: 10.1016/j.tvjl.2014.06.013. PubMed PMID: 25023088.

35. Glaser R, Harder J, Lange H, Bartels J, Christophers E, Schroder JM. Antimicrobial psoriasin (S100A7) protects human skin from Escherichia coli infection. Nat Immunol. 2005;6(1):57–64. Epub 2004/11/30. doi: 10.1038/ni1142. PubMed PMID: 15568027.

36. Elazar S, Gonen E, Livneh-Kol A, Rosenshine I, Shpigel NY. Essential role of neutrophils but not mammary alveolar macrophages in a murine model of acute Escherichia coli mastitis. Veterinary research. 2010;41(4):53. doi: 10.1051/vetres/2010025. PubMed PMID: 20416261; PubMed Central PMCID: PMCPMC2881416.

37. Gunn JS. Mechanisms of bacterial resistance and response to bile. Microbes Infect. 2000;2(8):907–13. Epub 2000/08/30. doi: 10.1016/s1286-4579(00)00392-0. PubMed PMID: 10962274.

38. Kandell RL, Bernstein C. BILE-SALT ACID INDUCTION OF DNA DAMAGE IN BACTERIAL AND MAMMALIAN-CELLS - IMPLICATIONS FOR COLON CANCER. Nutrition and Cancer-an International Journal. 1991;16(3-4):227–38. doi: 10.1080/01635589109514161. PubMed PMID: WOS:A1991GW53600008.

39. Bernstein C, Bernstein H, Payne CM, Beard SE, Schneider J. Bile salt activation of stress response promoters in Escherichia coli. Current Microbiology. 1999;39(2):68–72. doi: 10.1007/s002849900420. PubMed PMID: WOS:000081486100003.

40. Ahmadi S, Huang YC, Batchelor B, Koseoglu SS. BINDING OF HEAVY-METALS TO DERIVATIVES OF CHOLESTEROL AND SODIUM DODECYL-SULFATE. Journal of Environmental Engineering-Asce. 1995;121(9):645–52. doi: 10.1061/(asce)0733-9372(1995)121:9(645). PubMed PMID: WOS:A1995RQ38800008.

41. Iuliano A, Scafato P, Torchia R. Deoxycholic acid-based phosphites as chiral ligands in the enantioselective conjugate addition of dialkylzincs to cyclic enones: preparation of (-)-(R)-muscone. Tetrahedron-Asymmetry. 2004;15(16):2533–8. doi: 10.1016/j.tetasy.2004.07.009. PubMed PMID: WOS:000223666600012.

42. Huang WD, Hu TD, Peng Q, Soloway RD, Weng SF, Wu JG. EXAFS AND FTIR STUDIES ON THE BINDING OF DEOXYCHOLIC-ACID WITH COPPER AND ZINC IONS. Biospectroscopy. 1995;1(4):291–6. doi: 10.1002/bspy.350010407. PubMed PMID: WOS:A1995TG39100006.

43. Kokina TE, Salomatina OV, Popadyuk, II, Glinskaya LA, Korol’kov IV, Sheludyakova LA, et al. Complexes of Zn(II) and Cu(II) with the Amino Derivatives of Deoxycholic Acid: Syntheses, Structures, and Properties. Russian Journal of Coordination Chemistry. 2019;45(7):505–11. doi: 10.1134/s1070328419070030. PubMed PMID: WOS:000475588200006.

44. Hamner S, McInnerney K, Williamson K, Franklin MJ, Ford TE. Bile salts affect expression of Escherichia coli O157:H7 genes for virulence and iron acquisition, and promote growth under iron limiting conditions. PLoS ONE. 2013;8(9):e74647. Epub 2013/09/24. doi: 10.1371/journal.pone.0074647. PubMed PMID: 24058617; PubMed Central PMCID: PMCPMC3769235.

45. Ong CLY, Walker MJ, McEwan AG. Zinc disrupts central carbon metabolism and capsule biosynthesis in Streptococcus pyogenes. Scientific Reports. 2015;5. doi: 10.1038/srep10799. PubMed PMID: WOS:000355605300001.

46. Willis LM, Whitfield C. Structure, biosynthesis, and function of bacterial capsular polysaccharides synthesized by ABC transporter-dependent pathways. Carbohydr Res. 2013;378:35–44. doi: 10.1016/j.carres.2013.05.007. PubMed PMID: 23746650.

47. Russo TA, Liang Y, Cross AS. The presence of K54 capsular polysaccharide increases the pathogenicity of Escherichia coli in vivo. J Infect Dis. 1994;169(1):112–8. Epub 1994/01/01. doi: 10.1093/infdis/169.1.112. PubMed PMID: 8277173.

48. Russo T, Brown JJ, Jodush ST, Johnson JR. The O4 specific antigen moiety of lipopolysaccharide but not the K54 group 2 capsule is important for urovirulence of an extraintestinal isolate of Escherichia coli. Infect Immun. 1996;64(6):2343–8. Epub 1996/06/01. PubMed PMID: 8675348; PubMed Central PMCID: PMCPMC174077.

49. Buckles EL, Wang X, Lane MC, Lockatell CV, Johnson DE, Rasko DA, et al. Role of the K2 capsule in Escherichia coli urinary tract infection and serum resistance. J Infect Dis. 2009;199(11):1689–97. doi: 10.1086/598524. PubMed PMID: 19432551; PubMed Central PMCID: PMCPMC3872369.

50. Sarkar S, Ulett GC, Totsika M, Phan MD, Schembri MA. Role of capsule and O antigen in the virulence of uropathogenic Escherichia coli. PLoS ONE. 2014;9(4):e94786. doi: 10.1371/journal.pone.0094786. PubMed PMID: 24722484; PubMed Central PMCID: PMCPMC3983267.

51. Schneider G, Dobrindt U, Bruggemann H, Nagy G, Janke B, Blum-Oehler G, et al. The pathogenicity island-associated K15 capsule determinant exhibits a novel genetic structure and correlates with virulence in uropathogenic Escherichia coli strain 536. Infect Immun. 2004;72(10):5993–6001. doi: 10.1128/IAI.72.10.5993-6001.2004. PubMed PMID: 15385503; PubMed Central PMCID: PMCPMC517556.

52. Burns SM, Hull SI. Loss of resistance to ingestion and phagocytic killing by O(-) and K(-) mutants of a uropathogenic Escherichia coli O75:K5 strain. Infect Immun. 1999;67(8):3757–62. Epub 1999/07/23. PubMed PMID: 10417134; PubMed Central PMCID: PMCPMC96650.

53. Kim KS, Itabashi H, Gemski P, Sadoff J, Warren RL, Cross AS. The K1 capsule is the critical determinant in the development of Escherichia coli meningitis in the rat. J Clin Invest. 1992;90(3):897–905. Epub 1992/09/01. doi: 10.1172/JCI115965. PubMed PMID: 1326000; PubMed Central PMCID: PMCPMC329944.

54. Leying H, Suerbaum S, Kroll HP, Stahl D, Opferkuch W. The capsular polysaccharide is a major determinant of serum resistance in K-1-positive blood culture isolates of Escherichia coli. Infect Immun. 1990;58(1):222–7. Epub 1990/01/01. PubMed PMID: 2403532; PubMed Central PMCID: PMCPMC258433.

55. Bjorksten B, Bortolussi R, Gothefors L, Quie PG. Interaction of E. coli strains with human serum: lack of relationship to K1 antigen. J Pediatr. 1976;89(6):892–7. Epub 1976/12/01. doi: 10.1016/s0022-3476(76)80592-6. PubMed PMID: 792409.

56. Russo TA, Moffitt MC, Hammer CH, Frank MM. TnphoA-mediated disruption of K54 capsular polysaccharide genes in Escherichia coli confers serum sensitivity. Infect Immun. 1993;61(8):3578–82. Epub 1993/08/01. PubMed PMID: 8392976; PubMed Central PMCID: PMCPMC281046.

57. Anderson GG, Goller CC, Justice S, Hultgren SJ, Seed PC. Polysaccharide capsule and sialic acid-mediated regulation promote biofilm-like intracellular bacterial communities during cystitis. Infect Immun. 2010;78(3):963–75. Epub 2010/01/21. doi: 10.1128/IAI.00925-09. PubMed PMID: 20086090; PubMed Central PMCID: PMCPMC2825929.

58. Murphy KC, Campellone KG. Lambda Red-mediated recombinogenic engineering of enterohemorrhagic and enteropathogenic E-coli. Bmc Molecular Biology. 2003;4. doi: 10.1186/1471-2199-4-11. PubMed PMID: WOS:000188325600001.

59. Edwards RA, Keller LH, Schifferli DM. Improved allelic exchange vectors and their use to analyze 987P fimbria gene expression. Gene. 1998;207(2):149–57. doi: 10.1016/s0378-1119(97)00619-7. PubMed PMID: WOS:000072270400007.

60. Baym M, Kryazhimskiy S, Lieberman TD, Chung H, Desai MM, Kishony R. Inexpensive Multiplexed Library Preparation for Megabase-Sized Genomes. PLoS One. 2015;10(5). doi: 10.1371/journal.pone.0128036. PubMed PMID: WOS:000354931700100.

61. Zhou ZM, Alikhan NF, Mohamed K, Fan YL, Achtman M, Brown D, et al. The EnteroBase user’s guide, with case studies on Salmonella transmissions, Yersinia pestis phylogeny, and Escherichia core genomic diversity. Genome Research. 2020;30(1):138–52. doi: 10.1101/gr.251678.119. PubMed PMID: WOS:000506573000013.

62. Zhou ZM, Alikhan NF, Sergeant MJ, Luhmann N, Vaz C, Francisco AP, et al. GrapeTree: visualization of core genomic relationships among 100,000 bacterial pathogens. Genome Research. 2018;28(9):1395–404. doi: 10.1101/gr.232397.117. PubMed PMID: WOS:000443569600014.

63. Starr KF, Porsch EA, Heiss C, Black I, Azadi P, St Geme JW. Characterization of the Kingella kingae Polysaccharide Capsule and Exopolysaccharide. PLoS One. 2013;8(9). doi: 10.1371/journal.pone.0075409. PubMed PMID: WOS:000325423500066.

